# Molecular and structural innovations of the stator motor complex at the dawn of flagellar motility

**DOI:** 10.1101/2024.07.22.604496

**Authors:** Caroline Puente-Lelievre, Pietro Ridone, Jordan Douglas, Kaustubh Amritkar, Betül Kaçar, Matthew Baker, Nicholas Matzke

## Abstract

The rotation of the bacterial flagellum is powered by the MotAB stator complex, which converts ion flux into torque. The origin and evolution of this remarkable complex is understudied. Here, we perform the first phylogenetic and structural characterisation and classification of MotAB and nonflagellar relatives. Using 193 genomes sampled across 27 bacterial phyla, we estimated phylogenies and ancestral sequences, and generated AlphaFold predictions for all extant and reconstructed proteins. We then mapped them onto the phylogeny to determine patterns of diversity and distribution of structural innovations. We identify two discrete groups: the Flagellar Ion Transporters (FIT) and the Generic Ion Transporters (GIT). The FIT proteins are structurally conserved and have a square fold domain and a torque-generating interface (TGI). FIT proteins are divided into two clades, termed TGI4 and TGI5, referring to whether there have 4 or 5 short helices in the TGI. TGI5 motors are predominantly found in Proteobacteria and include the well-studied *E. coli* K12 system, while TGI4 motors are found in diverse phyla and include the Na^+^-powered polar motors of *Vibrio* (PomAB). The GIT proteins, on the other hand, are structurally diverse and lack these attributes. The interaction between the A and B subunits is conserved across the FIT and GIT proteins. The two subunits are jointly necessary for function, with the genes typically adjacent within an operon. Motility assays in *E. coli* show that the structural elements unique to FIT play an important role in flagellar motility. Our results indicate that the stator motor complex has a single origin and shares unique motility-related structural traits.

**Significance Statement:** Flagellar motility is a key feature in bacterial pathogenicity and survival. It allows bacteria to propel themselves and direct movement according to environmental conditions. We investigated the molecular and structural diversity of the stator motor proteins that provide the ion motive force to power flagellar rotation. This study integrates phylogenetics, 3D protein structure modeling, motility assays and ancestral state reconstruction (ASR) to provide insights into the structural mechanisms that first powered the flagellar motor. We provide the first phylogenetic and structural characterisation and classification of MotAB and relatives.

## Introduction

Bacterial flagellar motility is one of the most ancient forms of motility (1). It allows bacteria to propel themselves and direct their movement by rotating their flagella (2). Flagellar rotation is facilitated by the bacterial flagellar motor (BFM), a bidirectional rotary nanomachine powered by the ion motive force (IMF) harnessed by the stators (3). The stators are multimeric complexes that rotate the BFM by transducing the chemical energy from the ion flux into torque (4). Stators are made from an assembly of motor-associated transmembrane (TM) complexes and each consist of five A subunits (MotA for proton-powered flagella, PomA for sodium-powered flagella) and two B subunits (MotB and PomB, respectively) (5, 6).

The A and B subunits form an ion channel that crosses the periplasmic space and the cytoplasm. The ion-binding channel is formed by two A transmembrane regions (A-TM3 and A-TM4) with the TM region of the B subunit. The A-TM2 region acts as a membrane anchor of the cytoplasmic loop (4). The stator complex interacts with the motor to drive the rotation of the filament. This precise interaction occurs between the A subunit and FliG (residues R90/D288-D289 and E98/R281), and is highly conserved across bacterial lineages (4). This coupling results in conformational changes in the A subunit and activates force generation from ion transit across the inner membrane through these channels. This ion passage is modulated by a small domain adjacent to the TM region of the B subunit, known as the plug domain (7).

The evolutionary origins of the stator complex remain poorly understood. The widespread presence of the BFM across bacterial taxa, both gram-negative and gram-positive, suggests an early origin, possibly dating to the Last Universal Common Ancestor (8). MotAB share homology with other bacterial proton-coupled TM transport systems: ExbBD and TolQR from the TonB and the Tol-Pal systems respectively (9). These multi-protein complexes use IMF to energise active transport across the outer membrane and maintain membrane stability and integrity (10, 11). Previous phylogenetic work showed that ExbBD and MotAB form distinct monophyletic groups and share a common ancestry (8), but it remains unclear which, if any, resembles the progenital form. A scenario in which the last common ancestor of Bacteria was a Gram-negative diderm (2) would support the hypothesis that MotAB originated as an exaptation of pre-existing conformational variations of ExbBD/TolQR-like complexes to become the powerstroke mechanism in filament rotation (12). Determining the order by which structural innovations specific to each system occurred is possible by mapping protein structure traits onto a phylogeny. This approach can inform the evolutionary history of ancient proteins and provide a framework to characterise protein diversity based on their phylogenetic relationships and molecular and structural traits.

The aim of this study was to determine the phylogenetic relationships of the MotAB complex with its non-flagellar homologs, and to use this phylogenetic framework to systematically describe and characterise the structural diversity of the bacterial proton-coupled TM transport systems. To achieve this, we integrated deep homology searches, protein sequence alignments informed by protein structure and gene order, deep phylogenies using Bayesian inference, Ancestral Sequence Reconstruction (ASR), 3D structural predictions, and motility experimental assays.

## Results

### The A and B subunits are found as operons across bacterial genomes

Jackhmmer searches across the 193 sampled genomes recovered 746 potential homologs for MotA. Preliminary alignments were manually curated to exclude proteins with sequence similarity under 10% and outliers. Final alignments consisted of 379 proteins.

Homology searches for either the complete protein sequence or for the conserved transmembrane (TM) domain were less successful for the B subunits, reflecting low levels of sequence conservation. Proteins annotated as MotB or OmpA were successfully recovered, while well-known structural homologs such as ExbD and TolR (11) were not. Therefore, the gene immediately downstream of the *motA* homolog was used to build the B subunit data set. In all cases (except when the A subunit gene was duplicated), the B gene was found next to the A gene (Supplemental Data). The length of the MotA alignment was 295 residues, and 308 for MotB. The MotA subunit sequences show higher similarity amongst themselves with a mean pairwise identity of 17.7%, compared to the 10.6% for MotB.

### Flagellar stator complexes form a single clade

Congruence between the phylogenies of the A and B subunits was measured by estimating clade support entropy, a measure that describes the amount of uncertainty in a set of phylogenetic trees (13). BEAST 2 analyses recovered 3449 clades from the A dataset, with an entropy of 105.8, and 71518 clades from the B dataset, with an entropy of 256.7. Densitree visualisations confirmed that tree distribution in the B subunit tree is more diffused, particularly in the deeper branches of the tree with no statistical support. However, concatenating the A and B subunit datasets increased the phylogenetic information content (recovering only 1998 clades, with an entropy of 90.8) and the posterior probability values (Figure S1).

Two major clades can be identified in the resulting phylogeny: the first clade contains the MotAB complexes from *E. coli* K12 and other well-studied flagellar systems such as *Vibrio*, and the second clade includes its well-known structural homologs ExbBD and TolQR (11) (Figure 1). We term the group that contains *E. coli* MotAB the Bacterial Flagellar Ion Transporters (FIT) clade; and Bacterial Generic Ion Transporters (GIT) clade for the proteins that group with TolQR and ExbBD from *E. coli* (Figure 1). Of the 379 protein sequences included in the phylogenies, 107 belonged to the FIT clade and 272 to the GIT clade. The FIT proteins were further divided into two subclades: The first subclade includes *E. coli* MotAB, and the second one includes *Vibrio* PomAB. The MotAB clade includes only Gram-negative bacteria, predominantly Proteobacteria, but also Planctomycetota, Verrucomicrobia, Armatimonadetes, Gemmatimonadetes, Acidobacteria, and Nitrospirae. The PomAB clade comprises Gram-positive and Gram-negative bacteria from the following phyla: Aquificae, Proteobacteria, Bacillota, Spirochaetes, Planctomycetota, Acidobacteria, Deferribacteres, Chloroflexi, and Nitrospirae. The GIT clade shows several large clades made of complexes with unknown functions. Within the GIT clade, one subclade contains *E. coli* ExbBD and TolQR and proteins associated with gliding motility (AglRS) (14); and the second subclade includes a diverse set of proteins of unknown function.

**Figure 1.**
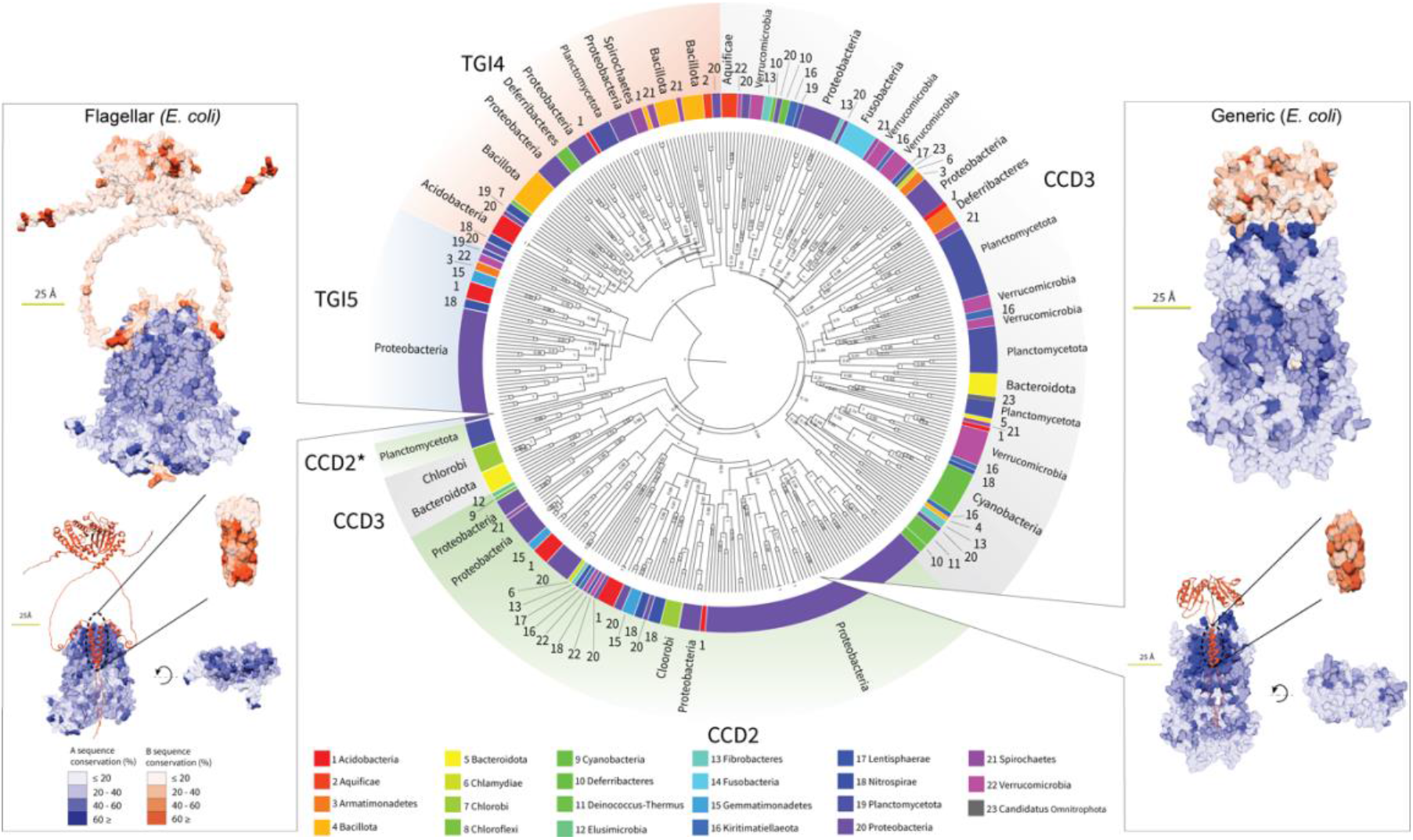
Taxonomic diversity mapped onto the AB concatenated phylogeny and comparison of residue conservation between the FIT and GIT A and B subunits. CCD-2* proteins form a separate clade from the rest of the CCD2 in the concatenated tree, but belong to the same clade in the A phylogeny. This likely reflects their idiosyncratic B subunit which has a unique N-terminal domain. CCD2: 2-helix Condensed Cytoplasmic Domain; CCD3: 3-helix Condensed Cytoplasmic Domain; TGI4: 4-helix Torque Generating Interface; TGI5: 5-helix Torque Generating Interface.

### Structural and molecular traits specific to the flagellar stators are conserved

AlphaFold predictions were generated for all 379 tips of the phylogenetic trees (Supplemental Data). Systematic comparison across all proteins showed the following structural traits are present in the FIT proteins and absent in the GIT proteins: 1) an Expanded Cytoplasmic Domain (ECD) with at least four helices that work as a Torque Generating Interface (TGI) in the A-subunit, 2) a four-helix structure in the shape of a square (the square fold) across the transmembrane layer in the A subunit, 3) a plug+linker domain in the periplasmic layer of the B subunit, and 4) an Expanded Peptidoglycan Binding (EPGB) domain in the periplasmic layer of the B subunit (Figures 1 and 4). The main structural difference between the two FIT subclades is the number of short helices in the Torque Generating Interface. One clade includes proteins with 4 short helices (TGI4), and the other clade includes proteins with 5 short helices (TGI5). The TGI4 subclade contains stator complexes with PomB-like plugs only, whereas MotB-like Plugs are found in both the TGI4 and the TGI5 subclades. Residue conservation in the PomB-like plugs is significantly lower than in the MotB-like Plugs (Figure S2). FIT proteins are conserved and display only 5 of 16 possible combinations of structural elements (Figure 2). Sequence conservation analyses show that the most conserved residues are those in the TM domains of the A and B subunits, which is where the interaction between the subunits occurs during ion flux (4) (Figure 1).

**Figure 2.**
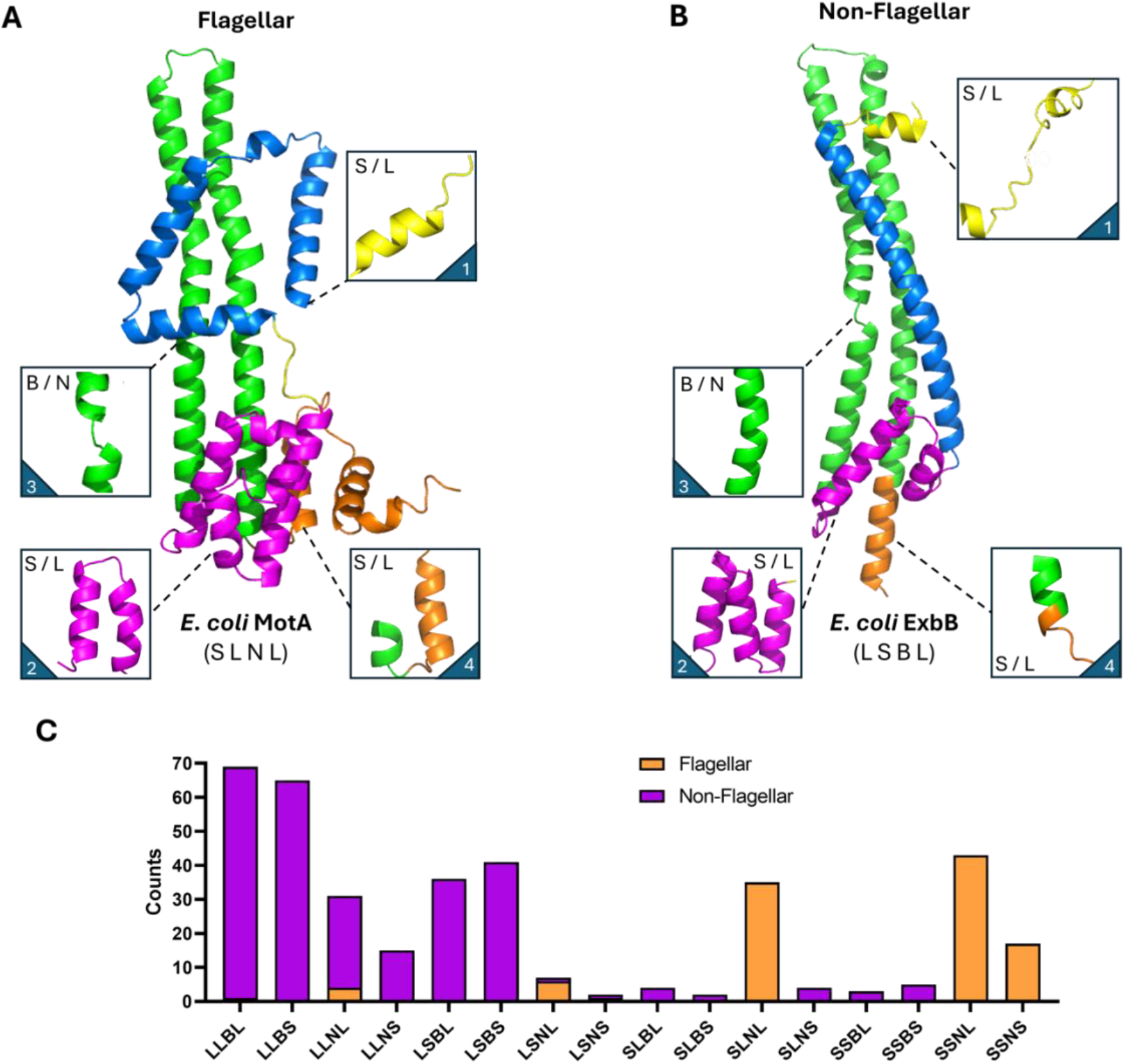
Structural Classification of A-subunit homologs. Monomeric models of (A) *E. coli* MotA and (B) *E. coli* ExbB coloured based on macroscopic structural features such as the N-terminal domain (yellow), TM domain (blue), the TGI domain (pink), TM3/4 (green) and C-terminal domain (orange). For each homolog, four regions of the protein, highlighted in insets, were considered for classification purposes: an N-terminal appendage (1, Small or Large), the TGI domain (2, Small or Large), TM3 (3, Broken or Not broken) and the C-terminal domain (4, Small or Large). Each homolog was expressed as a four-letter code as exemplified below each model (MotA: SLNL, ExbB: LSBL). (C) Bar chart summary of the classification for the whole A-subunit phylogeny. Entries belonging to the non-flagellar (purple) and flagellar (orange) clades were categorized and counted according to the 4-letter code shown on the x axis.

### High structural diversity across non-flagellar protein complexes

The GIT proteins show greater structural diversity, notably in their C and N-terminus elements of the A subunit. They also lack a Plug+Linker domain in the B subunits (Figures 1 and 3). The GIT proteins have a Condensed Cytoplasmic Domain (CCD), made of either two (CCD2) or three (CCD3) short helices. GIT proteins show 13 of 16 possible structural trait combinations (Figure 2) and in some cases, very large periplasmic domains of up to 305 residues (Supplemental Data).

**Figure 3.**
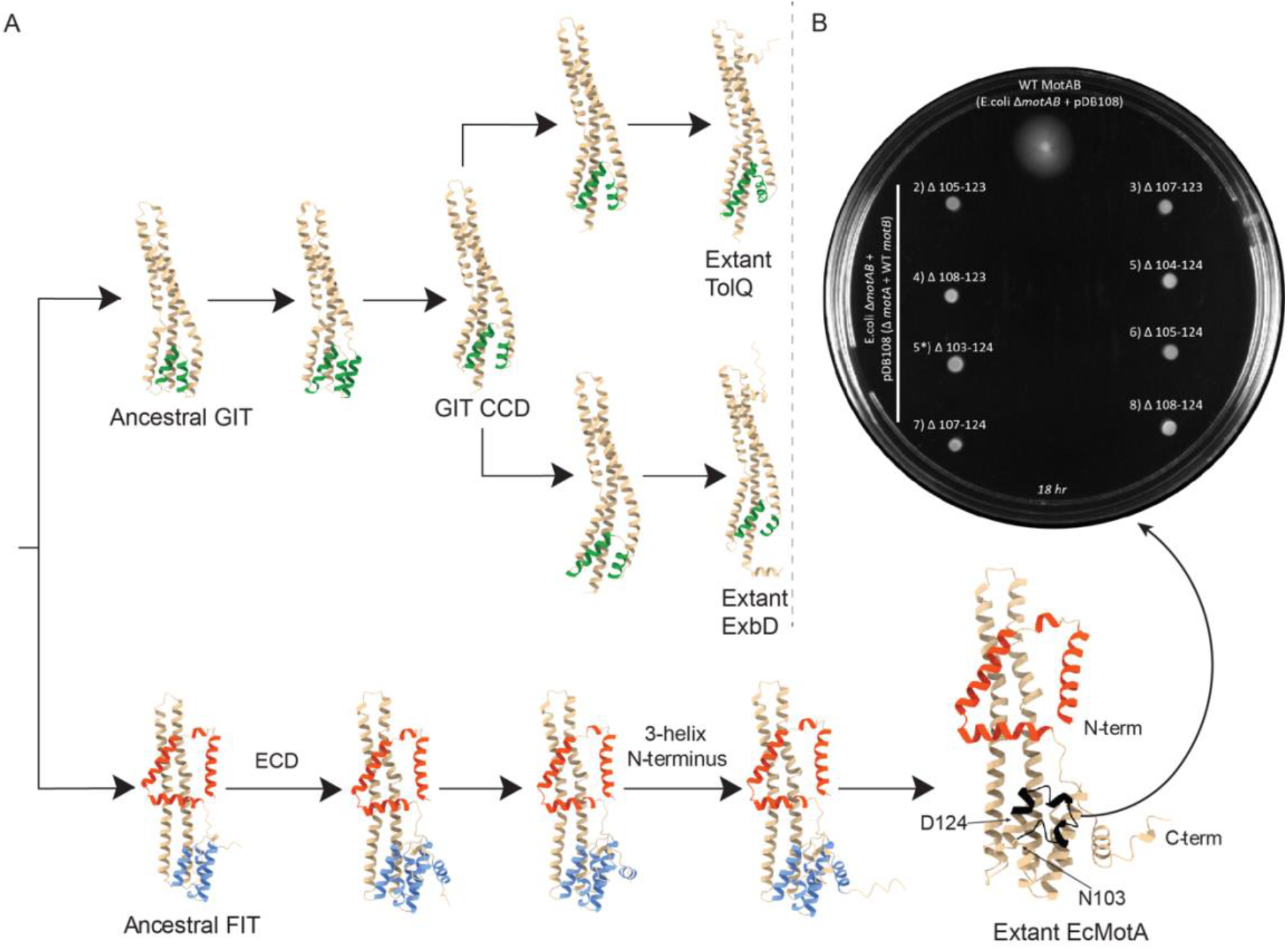
A) Hypothesised order of emergence of structural innovations in the Bacterial Flagellar Ion Transporters (FIT) and the Bacterial Generic Ion Transporters (GIT). Structure predictions for ancestral proteins are for the following nodes of the ASR phylogeny (Figure S4): 381, 543, 547, 548, 568, 652, 653, 657, 658. B) Soft-agar motility assay of TGI-deleted MotA variants in *E. coli* Δ*motAB*. Left) AF2 model of *Ec*MotA. The region ranging from N103 to D124 (black) of the TGI domain was targeted for deletions of various length. B) Soft-agar motility assay on 85mM Na^+^ LB agar (0.25%) plate (Ø=10cm) inoculated with *E. coli* Δ*motAB* expressing WT (top, motile) or deleted variants as labelled (2-9, non-motile). The plate was incubated for 18 hr at 30°C.

**Figure 4.**
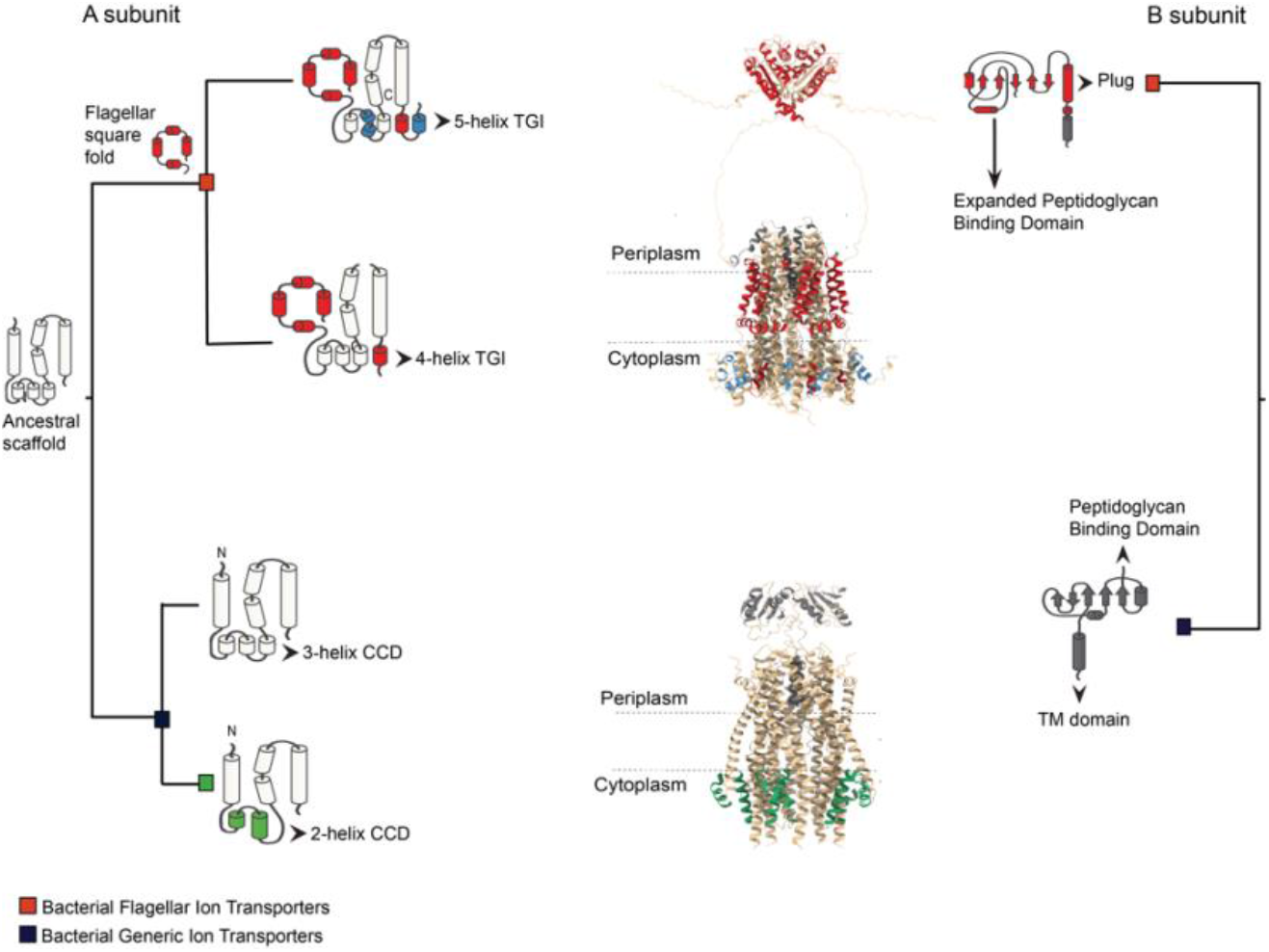
Simplified model of structure evolution of the A and B subunits of the bacterial ion transporters and the structural innovations associated with flagellar motility.

### Early structural divergence between FIT and GIT proteins

AlphaFold predictions of the reconstructed ancestral protein sequences suggest that the ancestor to FIT proteins had a square fold and a 3-helix CCD (Figure 3). Additional helices were gradually acquired and expanded the cytoplasmic domain to become TGI4 and TGI5. Conversely, the ancestral GIT protein appears to have had a linear transmembrane domain without a square fold. Unlike the FIT proteins, the CCD domain is further reduced to two helices in the CCD2 lineage (eg. ExbB and TolQ) (Figure 3). None of the structural elements that are characteristic of the extant FIT proteins was recovered in any of the nodes within the GIT clade.

### Removal of the TGI5 domain abolishes motility

Eight variants of *Ec*MotA with partially deleted TGI5 domains were generated and expressed. Deletions in MotA spanned residues N103–D124, a region of the protein encompassing the TGI5-specific structural elements as suggested by sequence alignments. Each of these variants was cloned in an arabinose-inducible, bicistronic plasmid (pDB108, pBAD33 backbone) together with WT *E. coli motB* and expressed in a *motAB* knock-out *E. coli* strain (*E. coli* Δ*motAB*), then tested on soft-agar swim plates for its ability to rescue motility and generate swim rings (Figure 3). All tested MotA variants were unable to rescue motility, while clones expressing WT *motAB* displayed swim rings typical of soft-agar motility assays (Supplementary Information). These results suggest that the TGI domain defined by residues N103-D124 is essential for motility in *E. coli*.

## Discussion

This study systematically surveyed and characterised the phylogenetic and structural diversity of the flagellar stator motor complex and their homologs across a broad range of bacterial lineages. Structural characterisation using modelling, sequence conservation analyses, and ASR suggests that the interaction between the A and B subunits is conserved across the FIT and GIT proteins. This interaction happens between the TM domains of both subunits, which seems to be homologous across all proteins. The neighbouring position of the A and B genes across all genomes sampled indicates that the two subunits typically work as an operon and are jointly necessary for function.

### Flagellar Ion Transporters

The FIT proteins showcase a square fold in their N-terminal domain, with the N-terminus pointing towards the cytoplasm. They do not exhibit any recognizable protein domain preceding the transmembrane, lipid facing, squared domain at their N-terminus (Figure 2). In their A subunit, the ECD is the point of contact with the rotor (FliG) and works as a TGI. The TGI can be made of 4 (TGI4) or 5 (TGI5) helices (Figures 2 and 4). Bacteria in the TGI5 clade are Gram-negative and predominantly Proteobacteria, including *E. coli*. Bacteria in the TGI4 clade belong to different lineages and include Gram-positive (Bacillota/Firmicutes) and Gram-negative (Figure 1). Motility assays in EcMotA showed that the lack of the TGI5 region results in the loss of motility in taxa that naturally have it. A possible explanation for this would be that the absence of TGI5 would alter the spatial configuration of the residues (i.e. R90) that are critical for the MotA-FliG interaction, resulting in loss of flagellar rotation. Alternatively, the TGI5 form could be involved in a different, potentially more specialised, type of regulatory mechanism of flagellar rotation.

The two well-defined FIT subclades (TGI4 and TGI5) could signal the existence of two different mechanisms by which the stator interacts with the rotor. Whether TGI4 and TGI5 interact differently with FliG or regulate rotation in different ways remains to be tested experimentally. Additional complete structures of rotor-stator complexes across TGI4 and TGI5 bacterial species will improve our understanding of the conserved rotor-stator interactions underlying flagellar rotation, how the different flagellar systems are regulated and their potential association or impact on bacterial lifestyle (15, 16).

### Generic Ion Transporters

The structural diversity of the GIT proteins is highlighted by a plethora of N-terminus appendages typical of the CCD2 clade that extend into the periplasmic space. The vast majority of them are domains of unknown function (DUF) eg. DUF3450 and DUF2341. It is possible that these appendages have evolved independently from the Exb/Tol-like transmembrane scaffold, and were later incorporated as proteins diverged and performed a broad range of biological functions. Future research focusing on characterising the functions of these DUF domains could help discover new biological systems and functions, such as the recently characterised Zorya anti-phage defence system (17). All GIT proteins lack a defined Plug+Linker domain and have a compact peptidoglycan-binding (PGB) domain in the B subunit. The low structure and sequence similarity between the GIT PGB domain and the bulkier EPGB of the FIT B subunits suggest that they are likely to have distinct evolutionary origins (11, 18).

### Final remarks

Our broadly sampled phylogeny of flagellar and non-flagellar MotAB homologs enabled us to map the distribution of structural traits and identify structural innovations unique to FIT complexes. We also discovered a major structural variation, the TGI5 domain, within the FITs that might be related to different mechanisms of flagellar motility regulation. We showed experimentally it is necessary for function in *E. coli*. A limitation of this study was exclusion of very remote homologs (such as the recently discovered ZorA and ZorB (17)) due to their very low sequence similarity (below 10%). This limitation may be overcome with new methods that incorporate structure directly into phylogenetic inference (19). Finally, our work suggests that using structural prediction to identify structural features diagnostic for well-supported major clades can inform a protein clade naming system that is more descriptive, stable and robust than ad hoc systems where names derived from experiments on a few model systems are inconsistently applied to related sequences.

## Materials and Methods

### Homology search and phylogenetic analyses

One hundred and ninety-three fully annotated and complete bacterial genomes were sampled across 27 phyla (Supplemental Data). When genome annotation was not available from NCBI (20), genomes were annotated using Prokka (21). The MotA sequence dataset was assembled using HMMER 3.3.2 (Nov 2020; http://hmmer.org/) to perform homology searches with MotA from *Escherichia coli* K12 as a reference (GenBank accession No. AAC74960.1). Jackhmmer was run for five iterations with the default search parameters. Since the genes for the A and B subunits are typically in synteny and function in an operon, the B subunits were identified with the gene immediately downstream from A. Identification of B subunits was confirmed manually with Genbank annotations and alignments. Protein sequences were initially aligned against a tailored HMMER profile that was generated from these datasets. Alignments were then refined with MAFFT (22, 23). Final alignments included only conserved domains. The A and B subunits were analysed separately and then concatenated for a partitioned analysis. Phylogenetic inference was performed using BEAST 2.7.6 (24). BEAST 2 analyses were conducted with the goal of estimating rooted trees, recognizing that without date calibrations the analysis produces only relative dating, rather than absolute dates. The analyses used an (uncalibrated) optimised relaxed clock (from the ORC 1.2.0 package (25)), the Yule skyline tree prior (BICEPS 1.1.2 (26)), and the OBAMA substitution model (OBAMA 1.1.1 (27)). The Yule skyline model assumes that diversification rates vary through time in a smooth piecewise fashion, providing a model-based estimate. The CCD0 tree was used to summarise the posterior distribution of trees (28). Tree entropy was measured to estimate phylogenetic information content in each posterior distribution of trees (13). Tree sets were visualised using DensiTree (29, 30). Phylogenetic trees were visualised using Figtree (http://tree.bio.ed.ac.uk/software/figtree/).

### Ancestral Sequence Reconstruction

ModelFinder (31) from IQ-Tree (32) was implemented to identify the best evolutionary model for the MotA sequence alignment (LG+F+R10). Ancestral sequence inference for the MotA phylogenetic tree was performed using Paml4.9 (33) (LG+F model). Sequence gaps were reconstructed by calculating the probability of a gap for each position based on a presence-absence sequence alignment and treating all positions with a gap probability >= 0.5 as a gap (34). Branch lengths were optimized based on the amino acid substitution models during ASR, to represent the expected number of amino acid substitutions per amino acid (33) (Figure S4).

### Sequence conservation analysis

Flagellar and non-flagellar subunits were distinguished based on the clade distribution in the concatenated phylogenetic tree and reference to experimentally characterized systems (PDB IDs 6YKM, 8GQY, 5SV0, 8ODT). Sequence alignments for the flagellar and non-flagellar MotA homologs were constructed using MAFFT with iteration refinement over 1000 cycles. Residues falling within a defined conservation range, as determined by sequence alignment, were allocated transparency levels using UCSF-Chimera (35).

### E. coli strains, plasmids, and culture media for swim assays

Stator variants were tested in stator-deleted derivative strains of *E. coli* RP437 (*E. coli* ΔmotAB) (36). The pDB108 (pBAD33 backbone, Cm+) plasmid encoding *motA* and *motB* (37) was used as the vector to clone and express all constructs. Deletions in the plasmid-encoded *motA* gene were generated using inverse-deletion PCR. Primers are listed in Table 1, Supplementary Information. PCR was used to linearize the pDB108 plasmid excluding nucleotides coding for the regions to be deleted. Linear PCR products were then re-ligated using T4 ligase (M0202S, New England Biolabs) and T4 Polynucleotide Kinase (Genesearch Pty Ltd) in T4 ligase buffer for 2 hr at 37°C followed by overnight incubation at 16°C. Every ligation reaction was then transformed into NEB10β competent cells (New England Biolabs). Plasmids were extracted from successful transformants for verification. Cloned constructs were confirmed by Sanger sequencing (Ramaciotti Centre for Genomics, University of New South Wales, Kensington, Sydney, Australia). Liquid cell culturing was done using LB broth (NaCl or KCl, 0.5% yeast extract (70161, Sigma-Aldrich), and 1% Bacto tryptone). Cells were cultured in agar plates composed of LB broth and 1% Bacto agar (BD Biosciences, USA). Swim plate cultures were grown in the same substrates adjusted for agar content (0.25% Bacto agar) and NaCL (85 mM).

### Structural modelling

AlphaFold predictions were generated for all the proteins included in the phylogenetic analyses and for the ancestral proteins estimated with ASR. Models of stator monomers and heptameric complexes were produced with AlphaFold2 (38) supported by the Australian AlphaFold Service (https://www.biocommons.org.au/alphafold) on the public server at UseGalaxy.org.au (39). The software was run in multimer mode using FASTA files as input (38, 40). Models were visualized using PyMOL version 2.5.4 (41).

### Quantification of structural diversity across A-subunit homologs

Alphafold2 monomeric models of all entries in the A-subunit phylogeny were manually inspected and classified as 4-letter codes according to the following principles: the first letter indicates the presence (L for large) or absence (S for small) of amino acid residues at the N terminus of the TM domain; the second letter indicates the TGI domain was composed of three or more α-helices (L) or two or fewer (S). The third letter indicates whether the TM3 helix was broken in two (B) or unbroken (N). The fourth and last letter indicated the presence of one (S) or more (L) α-helices at the C-terminus of the protein.

## Acknowledgments

This work was supported by the Human Frontier Science Program Grant No. RGY0072/2021. NJM was additionally supported by NZ RSTA grants 21-UOA-040 & 18-UOA-034 and BK was additionally supported by the John Templeton Foundation No 61926. JD is supported by the Alfred P. Sloan Foundation Matter-to-Life program Grant number G2021-16944. BK and KA thank the Center for High Throughput Computing at the UW-Madison. This work was supported by the Australian BioCommons which is enabled by NCRIS via Bioplatforms Australia funding. CPL thanks Diana Pese from the University of Auckland.

