## Supplemental tables and figures for "Molecular and structural innovations of the stator motor complex at the dawn of flagellar motility"

**Supplementary Information**

| *Primer Name* | *Sequence (5'→3')* |
|---|---|
| *MotA dTGI5 Fw-124* | *TTTATCGTCGATTATCTGCG* |
| *MotA dTGI5 Rv-104* | *ATTTTCAATATCACGTTCCAGC* |
| *MotA dTGI5 Rv-105* | *GGGATTTTCAATATCACGTTCC* |
| *MotA dTGI5 Rv-108* | *GCTCTCACGGGGATT* |

**Table 1.** Primers used for cloning truncated variants of *motA* in plasmid pDB108. Note that truncated construct MotA Δ6 (Δ103-124) was accidentally isolated while screening for successfully cloned constructs and was added to the panel of MotA truncations to test for motility.

| # | ID | Non-Flagellar | Flagellar |
|---|---|---|---|
| 1 | LLBL | 68 | 1 |
| 2 | LLBS | 65 | 0 |
| 3 | LLNL | 27 | 4 |
| 4 | LLNS | 15 | 0 |
| 5 | LSBL | 36 | 0 |
| 6 | LSBS | 41 | 0 |
| 7 | LSNL | 1 | 6 |
| 8 | LSNS | 1 | 1 |
| 9 | SLBL | 4 | 0 |
| 10 | SLBS | 2 | 0 |
| 11 | SLNL | 0 | 35 |
| 12 | SLNS | 4 | 0 |
| 13 | SSBL | 3 | 0 |
| 14 | SSBS | 5 | 0 |
| 15 | SSNL | 0 | 43 |
| 16 | SSNS | 0 | 17 |

**Table 2.** Structural classification of members of the A-subunit phylogeny according to the system described in Methods and represented as bar chart in Fig. 2C. Each member of the phylogeny, separated in Non-Flagellar and Flagellar homologs, is classified in one of the sixteen combination of structural elements, expressed as a 4-letter code (ID).

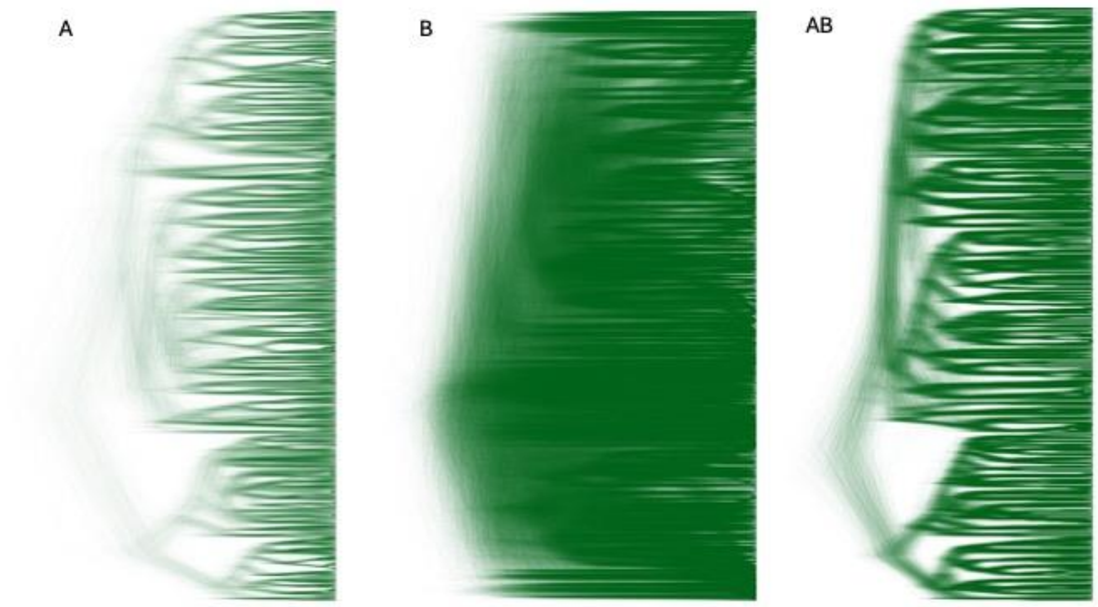


**Figure 1.** Densitrees for motA (A), motB (B) and motAB concatenated (AB).

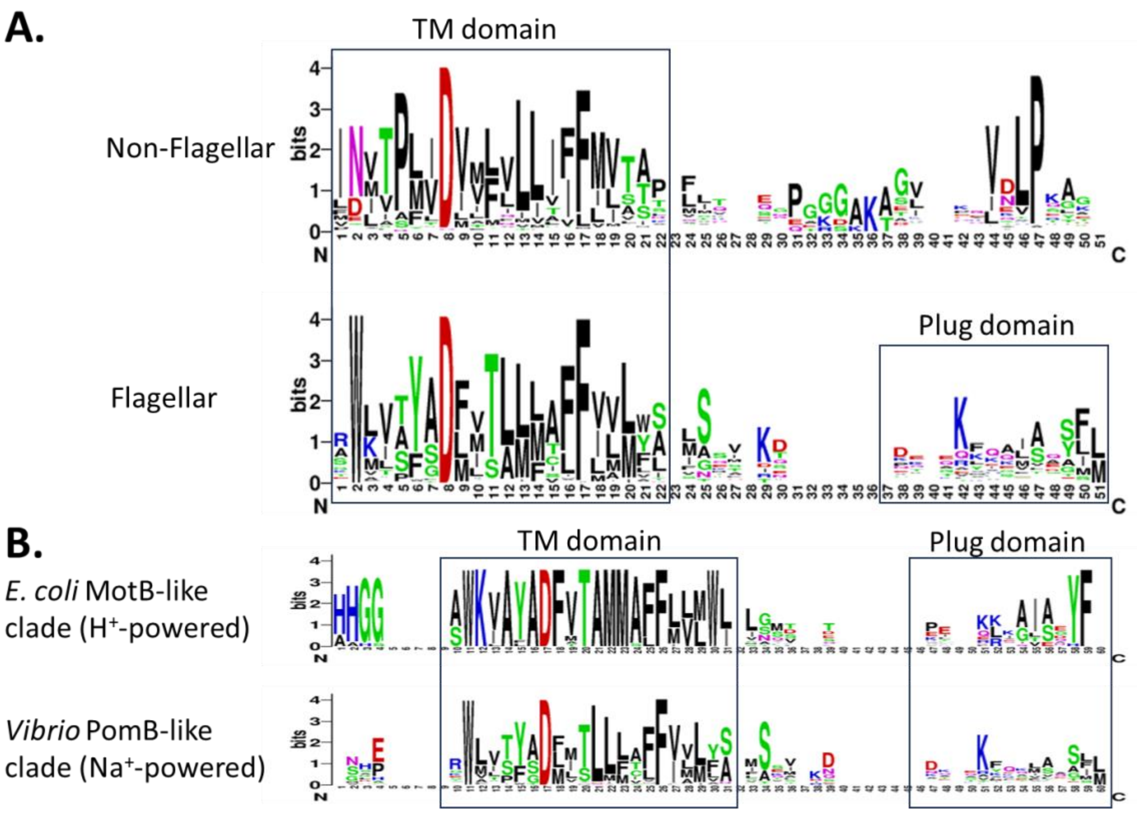


**Figure 2.** Residue conservation comparison between the FIT and GIT B subunits. A) Consensus logos are shown for the TM domain and Plug domain portions of B-subunit homologs belonging to Non-Flagellar (top) or Flagellar (bottom) clades. B) Consensus logos of the TM and Plug domains regions of homologs belonging to the same clade as the proton-powered *E. coli* MotB (top) or sodium-powered *Vibrio* PomB (bottom).

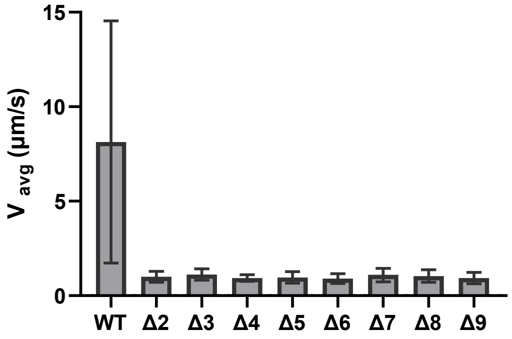


**Figure 3.** Average speed of free-swimming single cells in LB broth supplied with 1mM Arabinose, for wild-type and eight variants of *Ec*MotA with partially deleted TGI5 domains. Deletions spanned residues N103 – D124; WT MotA (N = 230 cells); Δ2: 105-123 (N = 333 cells); Δ3:107-123 (N = 267 cells); Δ4:108-123 (N = 148 cells); Δ5:104-124 (N = 295 cells); Δ6:103-124 (N = 324 cells); Δ7:105-124 (N = 320 cells); Δ8:107-124 (N = 318 cells); Δ9:108-124 (N = 353 cells). Error Bars indicate standard error of the mean. Statistical analysis was performed using pairwise Student's T test; **** P value < 0.0001.

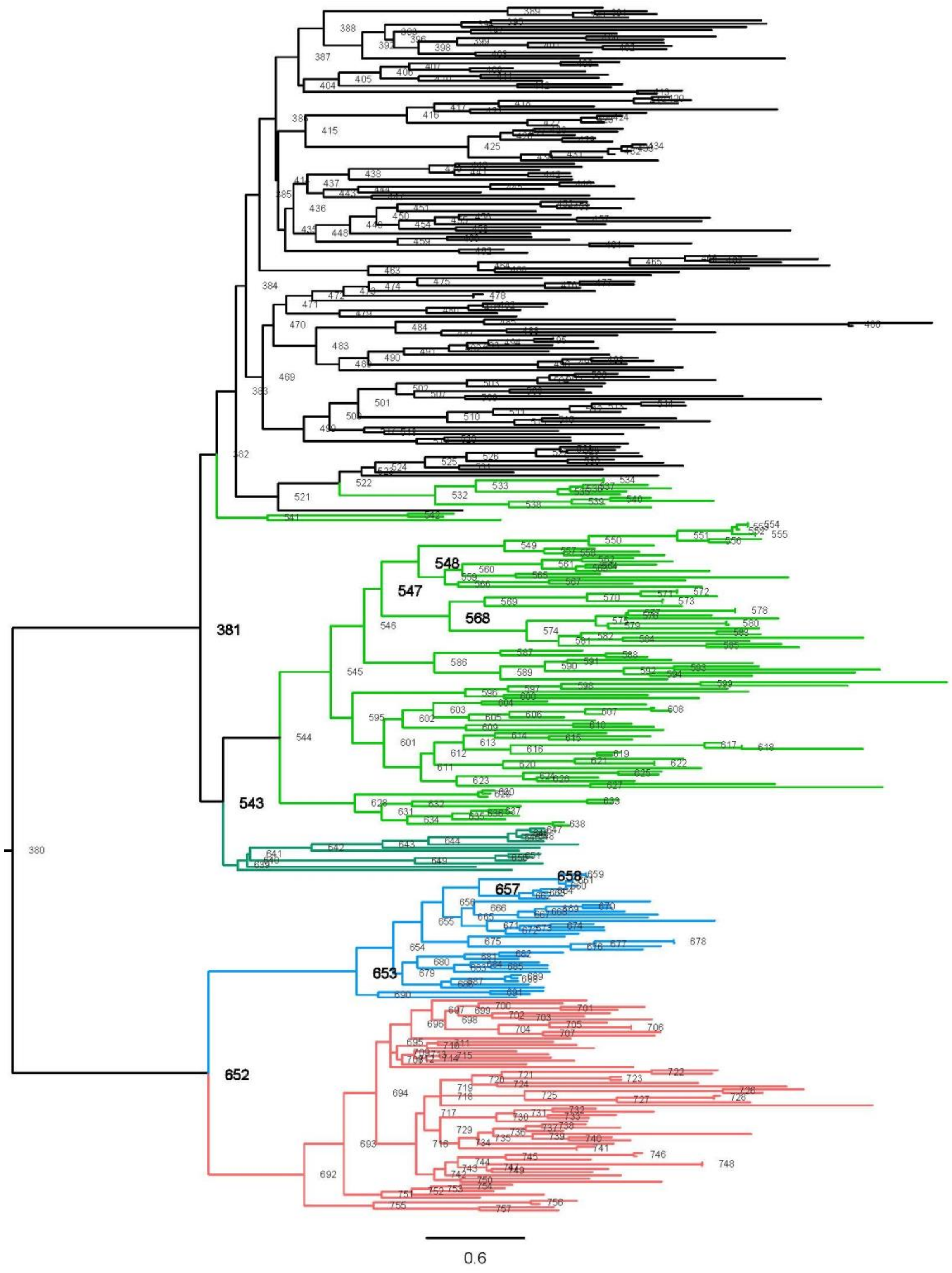


**Figure 4.** ASR phylogeny of the A subunits recoloured based on the largest structural features diagnostic of each clade. TGI4: red, TGI5: blue, CCD2*: dark green, CCD2: lighter green, CCD3: black. The nodes with structure predictions represented in Figure 3 are boldened in larger font.
